# Modeling and Analyzing Neural Signals with Phase Variability using the Fisher-Rao Registration

**DOI:** 10.1101/2020.04.30.069120

**Authors:** Weilong Zhao, Zishen Xu, Wen Li, Wei Wu

**Author notes:** Corresponding Author: Department of Statistics, Florida State University, 117 N Woodward Ave, Tallahassee, FL, 32306-4330, USA.

## Abstract

The Dynamic Time Warping (DTW) has recently been introduced to analyze neural signals such as EEG and fMRI where phase variability plays an important role in the data. In this study, we propose to adopt a more powerful method, referred to as the Fisher-Rao Registration (FRR), to study the phase variability. We systematically compare the FRR with the DTW in three aspects: 1) basic framework, 2) mathematical properties, and 3) computational efficiency. We show that the FRR has superior performance in all these aspects and the advantages are well illustrated with simulation examples. We then apply the FRR method to two real experimental recordings – one fMRI and one EEG data set. It is found the FRR method properly removes the phase variability in each set. Finally, we use the FRR framework to examine brain networks in these two data sets and the result demonstrates the effectiveness of the new method.

## 1 Introduction

Temporal phase lags have been commonly observed in neural signal recordings (Vinck et al., 2011; Williams et al., 2020). These lags may be introduced by the dynamic switching of brain states and exhibit linear or non-linearly patterns across the time domain (Allen et al., 2014; Chen et al., 2015). To make meaningful inference on the recordings, it is often a prerequisite to measure and remove those phase lags. For example, the study of functional connectivity focuses on identifying statistical interdependencies between time series recording from different brain areas (Friston et al., 1997; Tononi et al., 1998). One major goal is usually to quantify the strength of phase synchronization, which constitutes a significant physiological mechanism for functional integration (Fries, 2005; Singer, 1999; Singer and Gray, 1995; Varela et al., 2001). Various mathematical measures, such as phase-locking statistics (Lachaux et al., 1999), mutual information (Hurtado, 2004), and partial directed coherence (Kamiński et al., 2001), have been applied to quantify the synchronization. However, all these methods have clear limitations. For example, the phase-locking statistics is used to detect stationary, non-zero phase lags between two signals, whereas phase differences between different brain areas may vary dynamically, even with a single strong source. Similar limitation happens for the partial directed coherence method which is only appropriate for stationary and linear models.

In general, neurophysiological recordings often suffer from delay-induced bias. This includes those with high temporal resolutions such as electroencephalogram (EEG) and magnetoencephalography (MEG), and those with high spatial resolutions such as functional magnetic resonance imaging (fMRI). Indeed, small hemodynamic response delays or slice-timing differences may cause mismatches of data. To estimate the temporal latency, conventional approaches assume the delay-induced bias is stationary and shift signal curves within time scales of seconds to minutes. The estimated temporal delay is the time shift when strongest statistical interdependencies are obtained (Mørup et al., 2008). An increasing number of studies on fMRI (Allen et al., 2012; Chang and Glover, 2010; Handwerker et al., 2012; Jones et al., 2012; Kiviniemi et al., 2011; Sakoğlu et al., 2010; Smith, 2012), near-infrared spectroscopic and MEG (Brookes et al., 2011; de Pasquale et al., 2010; Keilholz, 2014; Li et al., 2015) indicate that functional connectivity reveals dynamic changes in human cognition process.

To capture a high-quality dynamic temporal-lag structure, various conventional approaches such as sliding-window analysis, wavelet transform coherence, and spontaneous co-activation patterns analysis, have been introduced to deal with this problem (Chen et al., 2015; Liu and Duyn, 2013). Traditional statistical methods such as linear and piecewise-linear models are also exploited (Williams et al., 2020). In particular, the dynamic time warping (DTW), a classical algorithm for measuring similarity between two temporal sequences (Sakoe and Chiba, 1978), has been a preferred method in recent studies of alignment on neural signals such as EEG (Gupta et al., 1996; Huang and Jansen, 1985; Karamzadeh et al., 2013), fMRI (Dinov et al., 2016; Meszlényi et al., 2017), and neuronal spike trains (Lawlor et al., 2018). Such temporal warping method offers a novel and enchanting insight to deal with common temporal-lag problems. The key idea in the DTW is to perform a dynamic time warping on discrete sequence data to correct the non-stationary time lags. The method is immune to the common noise components in the data and can provide accurate estimate of the underlying templates such as the hemodynamic response functions (Meszlényi et al., 2017).

Nonetheless, the DTW method has apparent disadvantages: 1) It is a matching method between two sequences of temporal points. The matching process on points are not one-to-one, and therefore the sequence lengths change after the alignment (cannot be pre-determined beforehand). 2) The matching does not preserve feature in the given sequences and often leads to the pinching effect problem (Srivastava et al., 2011). 3) The DTW is not a metric-based method, and can only do pairwise comparison. There is no principled way to conduct further statistical analysis (e.g. no notion of mean or covariance). 4) There is only pairwise comparison in the DTW method. It is very inefficient to use the method in practice. 5) Last but not least, the DTW measures the distance after the phase-lag is removed, and therefore this distance can be referred to as “amplitude distance”. However, the method does not provide a measurement on how much warping is done. That is, it does not provide a measurement on the “phase distance” between two signals. The detailed discussion of these disadvantages is given in Sec. 2.3.

To overcome these problems in the DTW, we propose to adopt a new framework, referred to as Fisher-Rao Registration (FRR), to deal with the dynamic latency between signals. Treating signals as functions on a given time domain, the FRR (Srivastava et al., 2011) offers a unified, comprehensive solution to conduct alignment (or registration) between two functions. Similar to the DTW, the alignment in the FRR is done with the notion of time warping, albeit as a continuous operation within the given domain. Any strictly increasing, nonlinear warping transformation is taken into account in this fixed domain. More importantly, the FRR is a metric-based method, and can naturally introduce the notion of mean and covariance on the given signals and produce more powerful statistical analysis. It can also provide both “amplitude distance” and “phase distance” between two signals. We will provide a systematical comparison between the FRR and the DTW and demonstrate the superiority of the FRR in this paper. We expect the FRR framework will turn out to be a useful method for neural signal processing where phase variability is a significant factor. The computational analysis in this study was implemented using Matlab (The Mathworks, Inc.), and scripts on examples can be accessed on GitHub ^1^.

The rest of this article is organized as follows. In Section 2, we will provide details of the FRR method and compare it with the DTW method in terms of basic framework, mathematical properties, and computational efficiency. In Section 3, we will illustrate the use of FRR method in real experimental data. We show that this method can effectively remove phase variability in given functional signals and such noise removal can be used to build more appropriate brain networks. Section 4 summaries the results and offers conclusions. Computational details on the DTW and FRR are given in Appendices.

## 2 Methods

### 2.1 Review of the Dynamic Time Warping method

The Dynamic Time Warping (DTW) (Sakoe and Chiba, 1978) is a finite, discrete, algorithm-based method that can match two curves, in the form of two finite-length sequences of sampled points, to correct the phase difference. The main idea is to minimize the distance between the two sequences through copying and matching the points. To achieve this, one needs to select one sequence as the base sequence, while the other one as the operating sequence, and define a sliding window. For each point in the operating sequence, we can go over all the points in the sliding window on the base sequence to compute the cost value. It is computed as the distance between the starting point to the current point on the operating sequence and the starting point to the matched point on the base sequence. In this way, after all cost values for the last point on the operating sequence are obtained, the minimum distance between the two sequences is defined as the cost of matching the last point on the operating sequence to the last point on the base sequence.

The implementation of the DTW follows a shortest-path-finding idea. Suppose 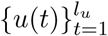 and 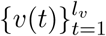 are two sequences with integer time index *t* and the sliding window has window size to be *w*, the cost *C*_*i,j*_ of matching *u*(*i*) to *v*(*j*), where *i* = 1, 2, …, *l*_*u*_ and *j* = max {1, *i* − *w*}, max {1, *i* − *w*}+ 1, …, min {*l*_*v*_, *i* + *w*}, is defined as: 

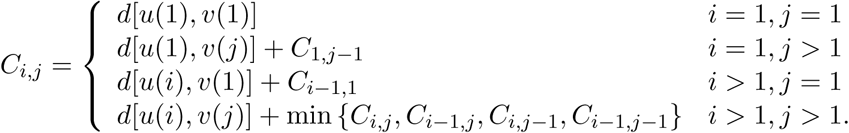

The *d*(*x, y*) in the formula is used to measure the squared distance between two points *x* and *y*. For example, when *x* and *y* are real numbers, a commonly used *d* will be the squared Euclidean distance: *d*(*x, y*) = (*x* − *y*)^2^. To compute the cost for matching *u*(*i*) to *v*(*j*), one needs to compute three adjacent “past” cost values *C*_*i*−1,*j*_, *C*_*i,j*−1_ and *C*_*i*−1,*j*−1_ first. In this way, the last cost 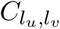 is obtained by an iterative way (loop over *i*, while for each *i*, loop over *j*). In each iteration, cost is computed and the three “past” costs given this cost are stored. Therefore, one can trace back from 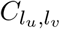 to get all previous matched costs one by one and figure out the shortest-path. The algorithm of the DTW method is given in detail in A.

A simple illustration of the DTW method is provided in Figure 1. Suppose there are two sequences of points which are sampled from two curves. Sequence 1 has 9 sampled points and Sequence 2 has 11 sampled points (see Figure 1(a)). The corresponding horizontal axis values are set to be just the index integers. The metric used is *d*(*x, y*) = (*x* − *y*)^2^. For simplicity, no sliding window is applied, i.e. *w* = 11. We need to note that the number of sampled points is different for the two curves and the aligned curves have more points than the original. The optimal path is shown in the matrix in Figure 1(b).

**Figure 1:**
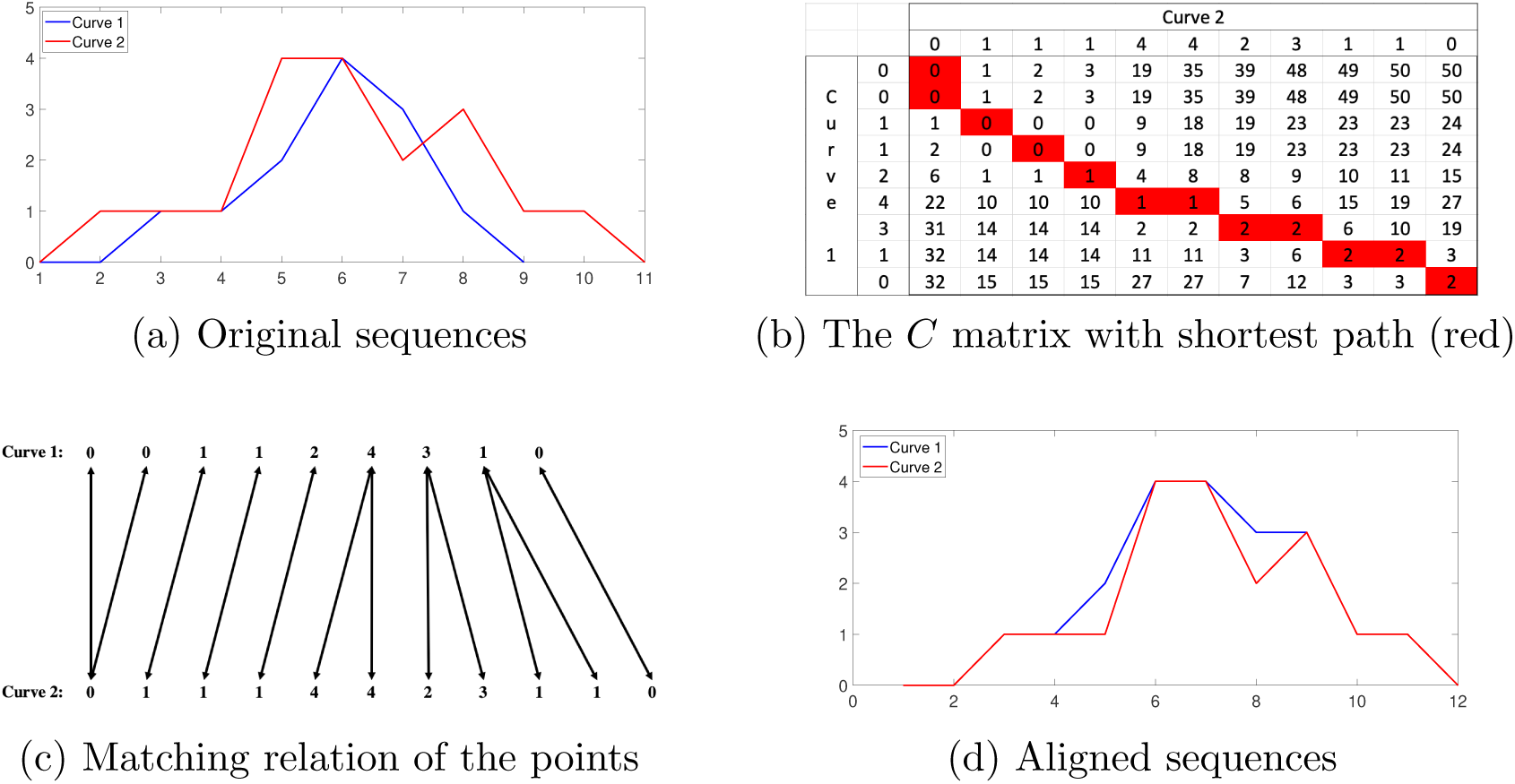
An illustration of the DTW method to match two curves in the form of two finite-length sequences of sampled points.

We note that in the framework of DTW method, the point matching is not one-to-one (see Figure 1(c)), which may corrupt global features in the original curves. The one-to-many matching also results in duplicates of some points and stretches the original curves by introducing more sampled points, shown in Figure 1(d). Moreover, since the DTW method is algorithm-based, it lacks solid mathematical background. One can only use finite, discrete numerical approximation to match two curves under the DTW method.

### 2.2 The Fisher-Rao Registration Method

#### 2.2.1 Basic Framework

The Fisher-Rao Registration (FRR) (Srivastava et al., 2011) is a nonparametric approach to conduct registration (or alignment) between two functions *f* and *g* on a time interval such as [0, 1]. The FRR method examines functions in the space *ℱ* = {*f* : [0, 1] → ℝ | *f* is absolutely continuous}. The alignment is represented by a time warping function in the set 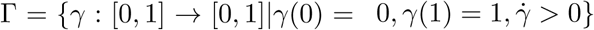, where 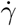 denotes the derivative function of *γ*. The set Γ is a *group* with identity *γ*_*id*_(*t*) = *t, t* ∈ [0, 1]. The main idea of the FRR is to find the optimal time warping function such that the extended Fisher-Rao distance between the two curves after warping is minimized, i.e. 

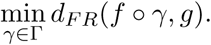

To achieve this, the concept of squared root velocity function (SRVF) is introduced:

##### Definition 1.

*For a function f* : [0, 1] → ℝ, *its SRVF is* 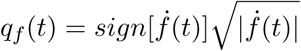

Given the SRVF function *q*(*t*) and function value at time 0, one can reconstruct the original function *f* by: 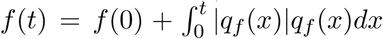. SRVF satisfies two important properties which build the registration framework:

##### Property 1.

*The Fisher-Rao distance between two functions is the same as the* 𝕃^2^ *distance between their SRVFs, i.e. d*_*F R*_(*f, g*) = ‖*q*_*f*_ − *q*_*g*_‖;

##### Property 2.

*For any function f and a time warping function γ, the SRVF of f ∘ γ is:* 

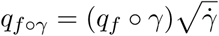

Based on the above two properties, the optimal time warping function between two functions *f* and *g* can be computed as: 

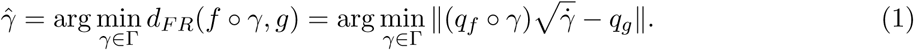

Computation of 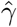 in Equation (1) can be efficiently done using a dynamic programming method (Srivastava et al., 2011).

Once the optimal warping function 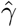 is known, there is no phase difference between *f ∘ γ* and *g*. The *amplitude distance* between *f* and *g* is given as: 

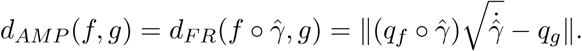

It is easy to verify that this amplitude distance is a semi-metric. That is, it satisfies nonnegativity, symmetry, and triangle inequality.

In addition to using 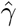 to measure the amplitude distance between *f* and *g*, we can use it to estimate the phase distance between them. Mathematically, we can measure the difference between 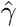 and *γ*_*id*_. Based on the Fisher-Rao metric, the *phase distance* is computed as: 

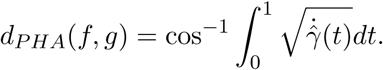

Note that the phase distance is in fact the arc length of *γ*_*id*_ and 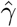 in their SRVF representations in the Hilbert unit sphere 𝕊^*∞*^. The distance is in the range [0, *π/*2].

In addition to the pairwise alignment for two functions, the FRR framework can also align a collection of functions. This is based on the fact that the amplitude distance is a semi-metric. In this case, we can define the notion of “mean” (or template) of the collection of functions and then align each function to the template. We briefly write down the estimation algorithm for the template of a collection of functions in B. The detailed procedure is given in (Srivastava et al., 2011).

### 2.3 Advantages of the FRR over the DTW

In this subsection, we will systematically compare the FRR and DTW methods and demonstrate advantages of the FRR. The comparison focuses on basic framework, mathematical properties, and computational efficiency.

#### 2.3.1 Basic Framework

As mentioned in Sec. 2.1, the DTW method is a point-to-point, finite, discrete method. In contrast, the FRR method is a function-based, continuous method. This difference makes the FRR method superior to the DTW method in two aspects: **domain for alignment** and **feature preservation**.

- **Domain for Alignment** The DTW method allows one-to-many and many-to-one mappings in order to obtain the minimal overall cost. Moreover, this method only focuses on the number of the sampled points, but not the exact time locations of these points. These properties often make the output of DTW method difficult to use, because the number of sample points will be changed in a non-systematical way after each alignment (the final number of points is case-by-case in each alignment and it cannot be known until after the alignment). In contrast, by considering the points as discrete observations on a curve, the FRR method naturally takes interpolation methods to make the number of points invariant during the alignment process. This difference is illustrated using the following example. As shown in Figure 2 (a), two curves: *f*_1_(*t*) = sin(*πt*) and *f*_2_(*t*) = 3 sin(*πt*) on [0, 1] are sampled at 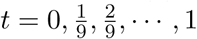 (10 points overall). It is clear that these two curves are already well aligned, so the ideal warping function between them should be the identity *γ*_*id*_(*t*) = *t*. This is what we obtain when the FRR alignment method is used and the result in shown in Figure 2 (b). In contrast, when the DTW method is applied on the 10 discrete points of the two curves, duplicate points are generated to minimize the cost. This is shown in Figure 2 (c). The number of sampled points increases from 10 to 14 after the alignment and the result is apparently not satisfactory. More importantly, there is no clear time locations for these 14 points in the warped curves.
- **Feature Preservation** In the DTW algorithm, the goal is to find the minimal sum of squared distance on the discrete points. The method may distort the given curves’ shapes for such purpose. This problem is well known as the *pinching effect* (Srivastava et al., 2011). This effect can corrupt the feature of the original curves by an improper alignment. On the contrary, the FRR method is based on a proper metric in the function space and can preserve features in the given functions. This comparison is illustrated with the following example. In Figure 3 (a), we generate two curves *f*_1_(*t*) = 5 sin(4*πt*^3^) and *f*_2_(*t*) = sin(4*πt*) on [0, 1]. The sampling points for *f*_1_ and *f*_2_ are both at 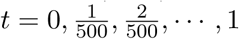 (501 points overall). It is observed that both curves have two peaks and two valleys, so an ideal mapping is supposed to result in peak-to-peak and valley-to-valley alignment. The alignment result using the DTW method is shown in Figure 3 (b). We can see that the DTW aligns the peak and valley, but destroys the original shapes entirely. On the contrary, the matching for FRR method not only aligns the peaks and valleys, but also keeps as much the original shapes as possible. This result is well demonstrated in Figure 3 (c) and (d).

**Figure 2:**
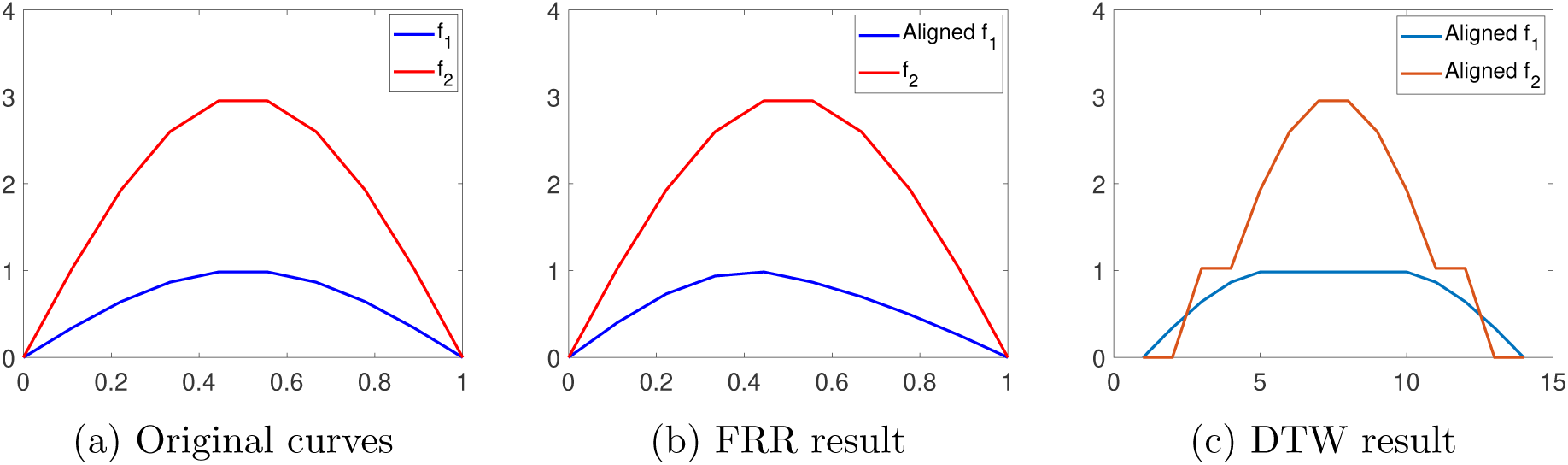
Illustration on the properness of the alignment. (a) Given two curves. (b) Alignment result using the FRR method. (c) Alignment result using the DTW method.

**Figure 3:**
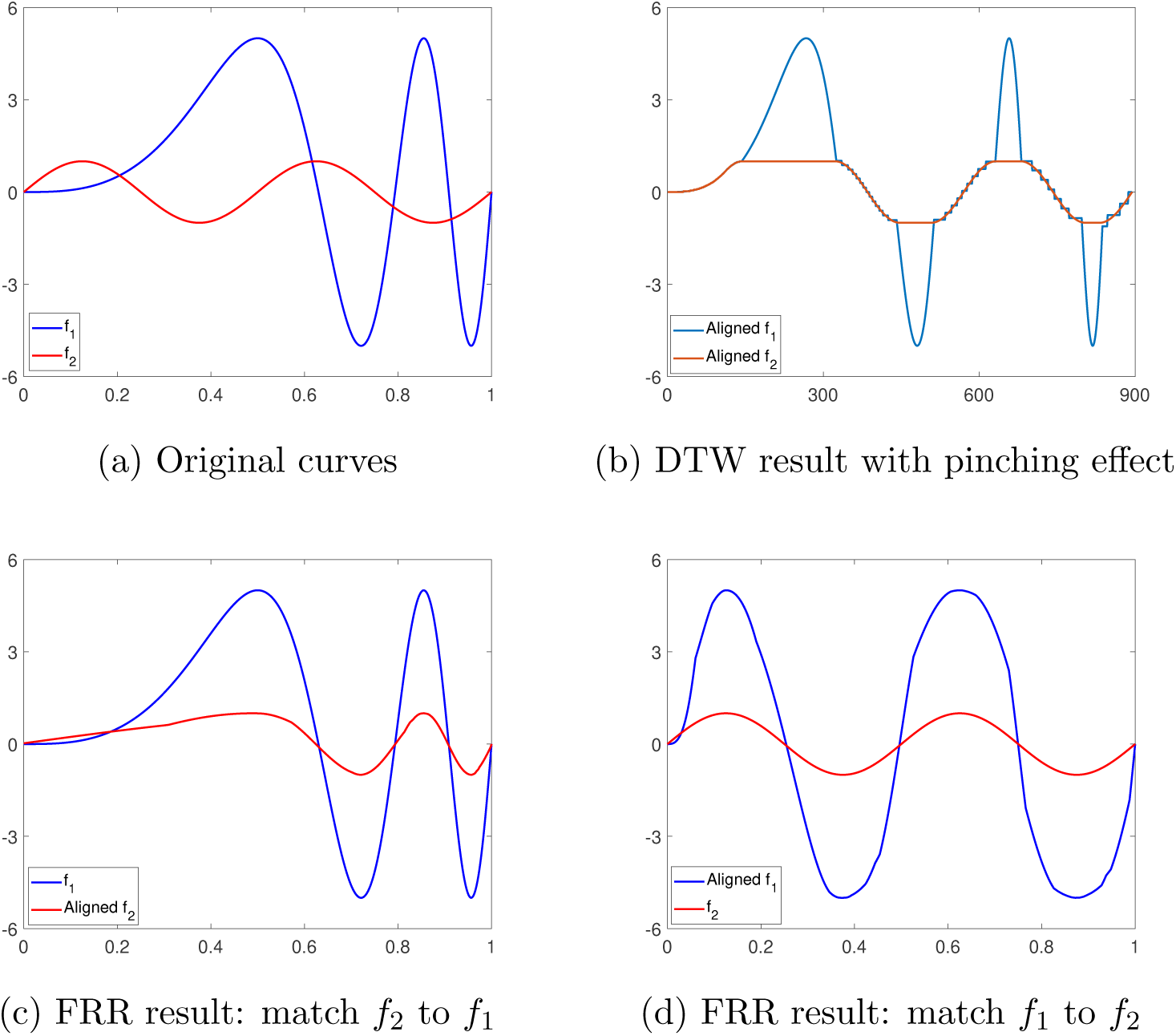
Illustration on the feature preservation. (a) Given two curves *f*_1_ and *f*_2_. (b) Alignment of *f*_1_ and *f*_2_ using the DTW method with a clear pinching effect. (c) Alignment of *f*_2_ to *f*_1_ using the FRR method. (d) Alignment of *f*_1_ to *f*_2_ using the FRR method.

#### 2.3.2 Mathematical Properties

As we have pointed out, the DTW is a procedure-based method on discrete sequences. The cost function does not present any metric distance between two curves. In contrast, the FRR follows a rigorous mathematical framework. The optimal distance between two curves naturally defines an amplitude metric and a phase metric. Based on these properties, we can introduce the notion of mean and covariance, and produce more powerful statistical analysis tools such as functional principal component analysis.

- **Metric property** As mentioned in the previous sections, the FRR method is derived from the Fisher-Rao distance. As a result, its framework is metric-based and the alignment is a matching of two curves. In contrast, the DTW method aims at minimizing the cumulative point-to-point distance and is not metric-based. One advantage of the metric-based method is that the matching relation can be well captured using the warping functions. We can learn how one curve is warped to another in the given time domain and measure the degree of warping in the alignment process (using the phase distance *d*_*PHA*_). On the contrary, the non-metric-based DTW can only obtain the point-to-point relation, and the mapping will change both of the curves to achieve the points’ alignment. We will use Figure 3 again to illustrate this difference. Basically, Figure 3 (a) shows the original curves *f*_1_ and *f*_2_, where the peaks and valleys present at different time locations. By the DTW algorithm, mapping from *f*_1_ to *f*_2_ has the same result as that from *f*_1_ to *f*_2_. This alignment result is shown in Figure 3 (b). In this case, both sequences change length after the alignment (increasing from 501 to to 900). Using Equation (1) in the FRR method, we can estimate the optimal warping function from one function to an objective function where the objective one remains unchanged. This is shown in Figure 3 (c) and Figure 3 (d). We can see that matching *f*_1_ to *f*_2_ and vice versa are two opposite transformations.
- **Statistical analysis on multiple observations** When dealing with multiple-curve alignment, the metric-based FRR method can result in more powerful analysis tools. Based on the metric distance, we can naturally define the mean of a set of functions 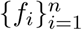 as 

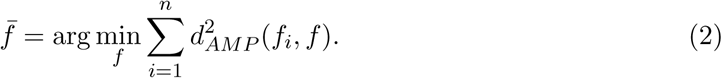 As the objective function can remain invariant, we can align every function to the mean (or the mapping template), and then all functions can be aligned altogether. Higher order-statistics such as covariance can also be computed using the aligned curves. Moreover, using the phase distance, we can measure the degree of phase change of each curve from the template. Further analysis tools can be adopted on the amplitude and phase result from the FRR method. For example, one can do functional principle component analysis (fPCA) to learn the detailed structure of the given curves (Tucker et al., 2013). One can also do functional analysis of variance (fANOVA (Zhang, 2013)) to compare different groups of curves. All these analysis methods are not available to the DTW method because it is not a metric-based method, where the pairwise alignment is the only result one can get. We here use one example to illustrate the FRR-based functional PCA. Let 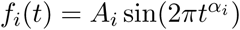, *i* = 1, 2, …, 10, *t* ∈ [0, 1], where *A*_*i*_ and *α*_*i*_ are independently and uniformly distributed with *A*_*i*_ ∼ *U* (1, 5) and *α*_*i*_ ∼ *U* (1, 3). All the ten curves are shown in Figure 4 (a). It is apparent that *A*_*i*_ determines the amplitude variation and *α*_*i*_ determines the phase variation for each curve. We at first examine functional PCA on the given 10 curves and the result is shown in Figure 4 (d). We show the mean curve, 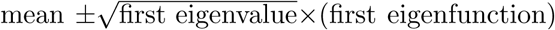 and 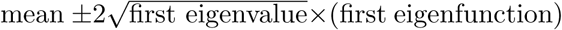. It is found that the first principle component obtained by applying functional PCA directly only explains around 65% of the total variance. That is, the first principal direction can only partially capture the variability in the given data. As a comparison, we conduct phase-amplitude separation for the original curves under the FRR framework, and then do functional PCA on amplitude components and phase components separately. The alignment results on aligned curves and time warping functions are shown in Figure 4 (b) and (c), respectively. Here, the aligned curves represent the amplitude components and the warping functions represent the phase components. Functional PCA on aligned curves and warping functions are shown in Figure 4 (e) and (f), respectively. It is found that the first principle component for aligned curves explains about 99% of total amplitude variance and the first principle component for warping functions explains about 97% of total phase variance. That is, the variability in amplitude and phase can nearly entirely captured by the first principal direction, respectively. These results show that for functional observations with phase variability the FRR method on amplitude and phase component provides a better structure representation than that on the original curves. In contrast, the DTW method is not able to do any modeling analysis.

**Figure 4:**
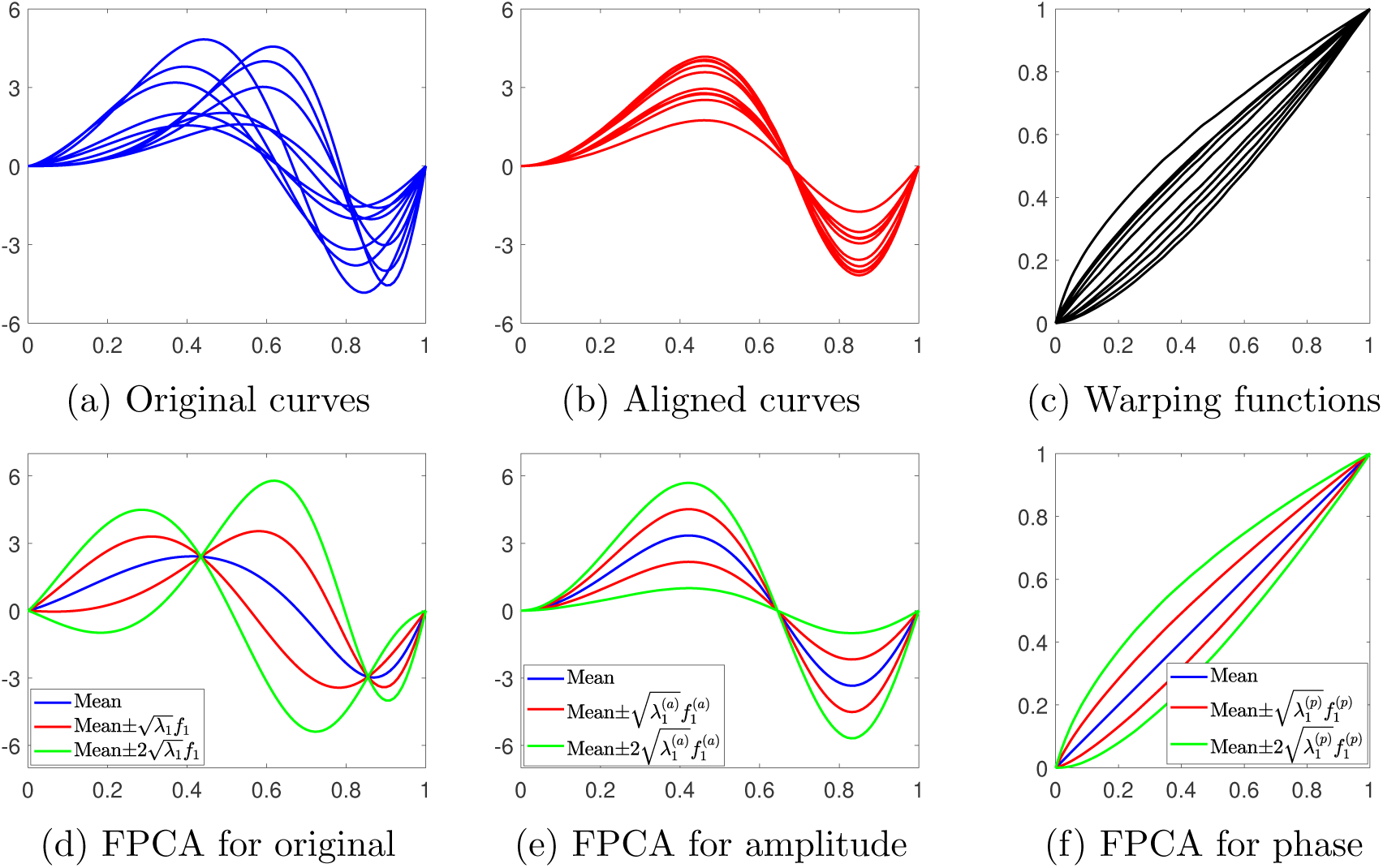
Illustration on fPCA with FRR framework. **(a)** 10 simulated curves; **(b)** Aligned curves obtained via the FRR method; **(c)** Time warping functions obtained via the FRR method. **(d)** FPCA result on the original curves: blue line represents the mean curve, red curves represent mean 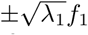, and red curves represent mean 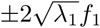, where *λ*_1_ denotes the first eigenvalue and *f*_1_ denotes the first eigenfunction in the original data; **(e)**: Same as (d) except for the aligned curves, where 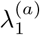 denotes the first eigenvalue and 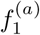 denotes the first eigenfunction in the aligned curves. **(f)**: Same as (d) except for the warping functions, where 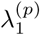 denotes the first eigenvalue and 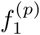 denotes the first eigenfunction in the warping functions.

#### 2.3.3 Computational Cost

As mentioned in Section 2.2.1, 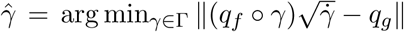 can be numerically solved using a dynamic programming procedure. This is the same framework as the DTW, and therefore for pairwise alignment the FRR and DTW have the same computational efficiency. However, as the FRR can naturally introduce summary statistics, it will have more efficiency in practical use for a large sample of data.

For example, assume we have two groups of curves 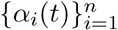 and 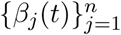, where the main difference between these two groups is the amplitude (the phase variability is treated as noise). We then observe unlabeled new curves 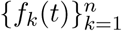 and our goal is to classify these new curves to one of the two classes using the nearest neighborhood (NN) method.

For the DTW method, the minimum cost between each pair of function is naturally taken as the “distance” in the NN method. For each new curve, we need to calculate such DTW distance between it and every curve in 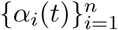 and 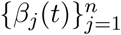. These pairwise computations are in the order 𝒪(*n*). Therefore, the total cost for all testing curves are in the order 𝒪(*n*^2^).

In contrast, using the FRR method, the *mean* curve of each group can be efficiently estimated in the computational order 𝒪(*n*) (Srivastava et al., 2011). For each new observation, we only need to compute the FRR distance to the two means, and then assign its label based on the smaller distance. Therefore, for all *n* test curves, the total computational cost is in the order of 𝒪(*n*) only.

This is apparently a significant advantage over the DTW method.

Finally, Table 1 provides a summary of all above comparisons of the DTW and FRR.

**Table 1:** Summary of comparisons between the DTW and FRR

| Method |  | DTW | FRR |
| --- | --- | --- | --- |
| Basic Framework | Domain | discrete sequence, length varied after alignment | continuous curve, length-invariant |
|  | Features | features distorted (pinching effect) | features well preserved |
| Mathematical Properties | Metric Property | not metric-based | metric-based |
|  | In-depth Analysis | pairwise distance only, no statistical analysis | summary statistics, fPCA, fANOVA, etc |
| Computation Cost in Group Classification | | pairwise only, $\mathcal{O}(n^2)$ | statistics-based, $\mathcal{O}(n)$ |

## 3 Application in Neural Signal Analysis

In this section, we apply the FRR method to neural signal analysis on an fMRI and an EEG data set. We will illustrate the FRR method using sample signals, and then use the FRR method to examine brain networks.

### 3.1 fMRI and EEG Data

We will at first describe the two datasets we use in this study.

#### 3.1.1 Data description

The fMRI data set was taken from Addiction Connectome Preprocessed Initiative ^2^. In particular, we use a subset of rest-fMRI data in the project of Multimodal Treatment of Attention Deficit Hyperactivity Disorder, which is per-processed by the National Institute on Drug Abuse (ADHD). The ADHD subset we use consists of 60 subjects, with 30 non-marijuana users and 30 marijuana users. The 6-minute resting-state fMRI were band-pass filtered (0.01Hz to 0.1Hz) and 116 brain regions-of-interest (ROIs) are drawn using the anatomical labelling atlas (Craddock et al., 2012), which include precentral gyrus, olfactory cortex, hippocampus and other areas. The preprocessing procedure results in 180 total time points at 0.5 Hz temporal resolution for each subject. Figure 5 shows preprocessed resting-state fMRI functional timeseries of two example ROIs in the marijuana-user group.

**Figure 5:**
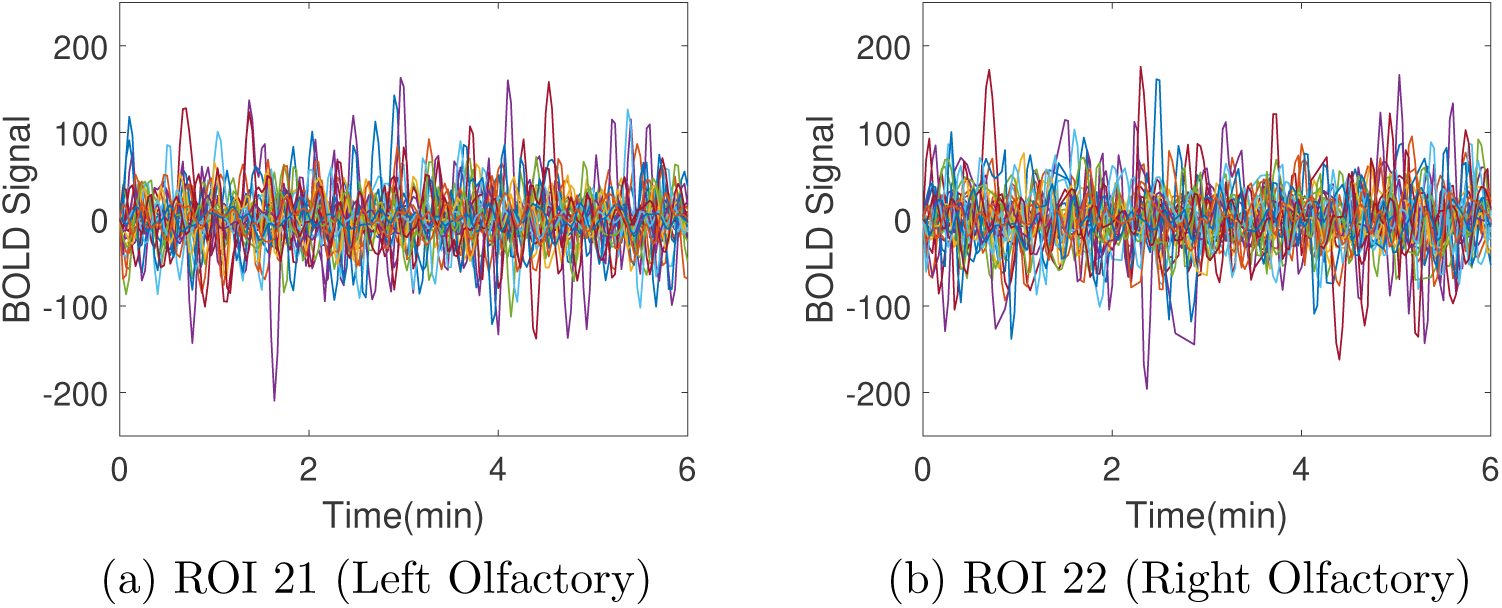
fMRI data in two example brain regions of 30 marijuana users. **(a)** Left olfactory. **(b)** Right olfactory.

EEG data were recorded at the Florida State University as a part of a prospective study (Clancy et al., 2018), where participants undertook four days of transcranial alternating current stimulation (tACS) at the alpha frequency (8-12 Hz), and resting-state EEG (2 minutes long) was recorded immediately before (baseline), immediately after, and 30 minutes after tACS on Day 1 and Day 4. The raw data was collected from 32 scalp electrodes of 33 subjects’ brains at 250 Hz temporal resolution. The scalp map for this experiment is shown in Figure 6(a). The baseline EEG recording before tACS on Day 1 and Day 4 are taken in this study. The EEG signals were then bandpass (8-12 Hz) filtered to contain alpha oscillations. EEG data signals in two example electrodes (AF3 and AF7) in Day 1 are shown in Figure 6(b)(c).

**Figure 6:**
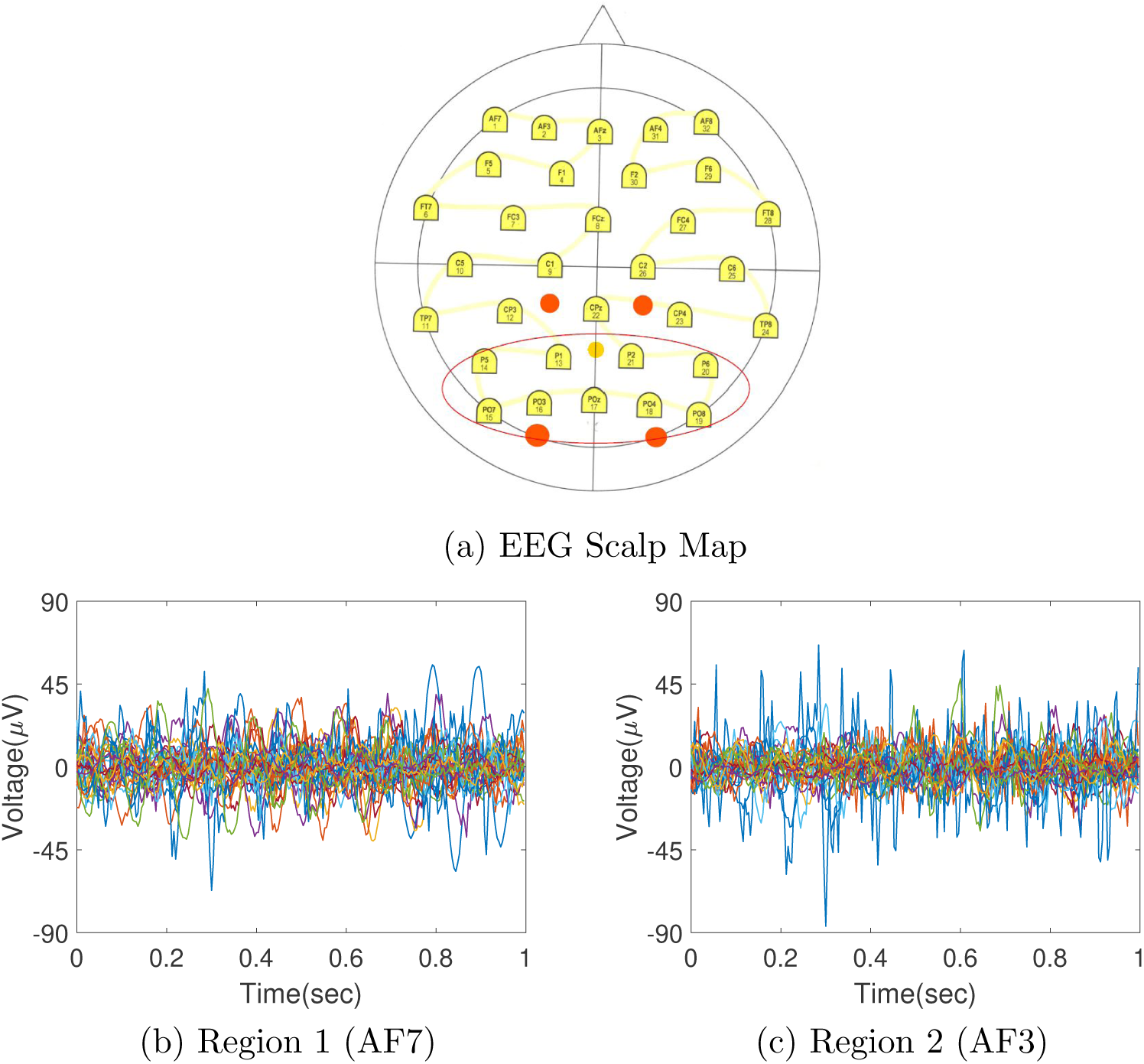
EEG data. **(a)** EEG Scalp Map (Clancy et al., 2018). **(b)** Data in the brain region AF7 of 33 subjects. **(c)** Same as (b) except for region AF3.

#### 3.1.2 Illustration of Alignment

For illustrative purposes, we compare the alignment results by the FRR and DTW methods using samples from the fMRI and EEG data sets described above. In particular, we select two typical timeseries from all 30 marijuana-users from the same ROI (Left Precentral) in the fMRI set. Similarly, we select two typical timeseries from Electrode 1 (AF7) of the Day 1 group in the EEG set.

The alignment result using the FRR method is shown in Figure 7. In Panel (a), we show two original fMRI signals with time length 3 minutes, denoted as *f*_1_ to *f*_2_. These two signals have apparent nonlinear phase lag between them. We then adopt the FRR framework to conduct alignment. The aligned result is shown in Panel (b), where we align *f*_2_ to match *f*_1_. We can see that all peaks and valleys are well matched after the alignment process. The optimal warping function is shown in Panel (c), which is close to the identical time warping function *γ*_*id*_(*t*) = *t*. Similar alignment on a pair of EEG signals with a length of 1 second is shown in Panels (d), (e), and (f). We can see that the FRR method can also align peaks and valleys in the signals.

**Figure 7:**
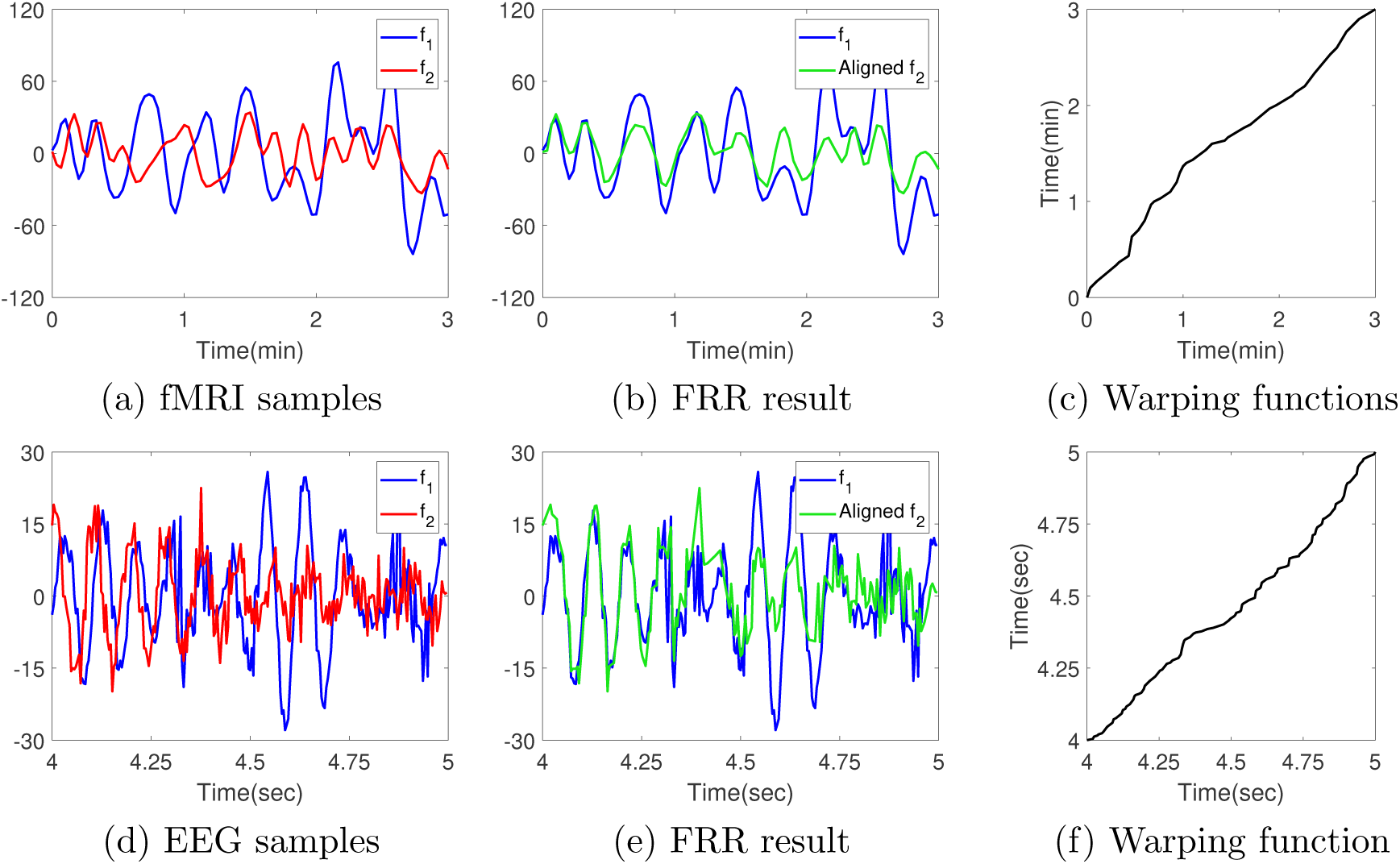
Illustration of the FRR method on real data. **(a)** One pair of fMRI signals *f*_1_ and *f*_2_; **(b)** *f*_1_ and the aligned *f*_2_ *∘ γ*; **(c)** Optimal time warping function; **(d), (e), (f)** Same as (a), (b) and (c) except for a pair of EEG signals.

In contrast, we evaluate the performance of the DTW method on the same fMRI and EEG signals and the result is shown in Figure 8. Note that the DTW method is constructed via aggregating point-wise deviations for discrete data, and the length of discrete observations are not consistent before and after DTW “alignment”. In Panel (a), we show the observed sequences with 90 discrete sampling points. The aligned result is shown in Panel (b), where pinching effect is clearly observed and the number of points increase to 120. Instead of aligning peaks and valleys, a lot of flat segments are generated which corrupt the basic features in the original data. This pinching effect is supported by the shortest path in Panel (c). We see several horizontal and vertical segments which indicates one-to-many matching. Similar alignment on a pair of EEG signals with 250 sampling points is shown in Panels (d), (e), and (f). We can also see the pinching effect in the alignment result.

**Figure 8:**
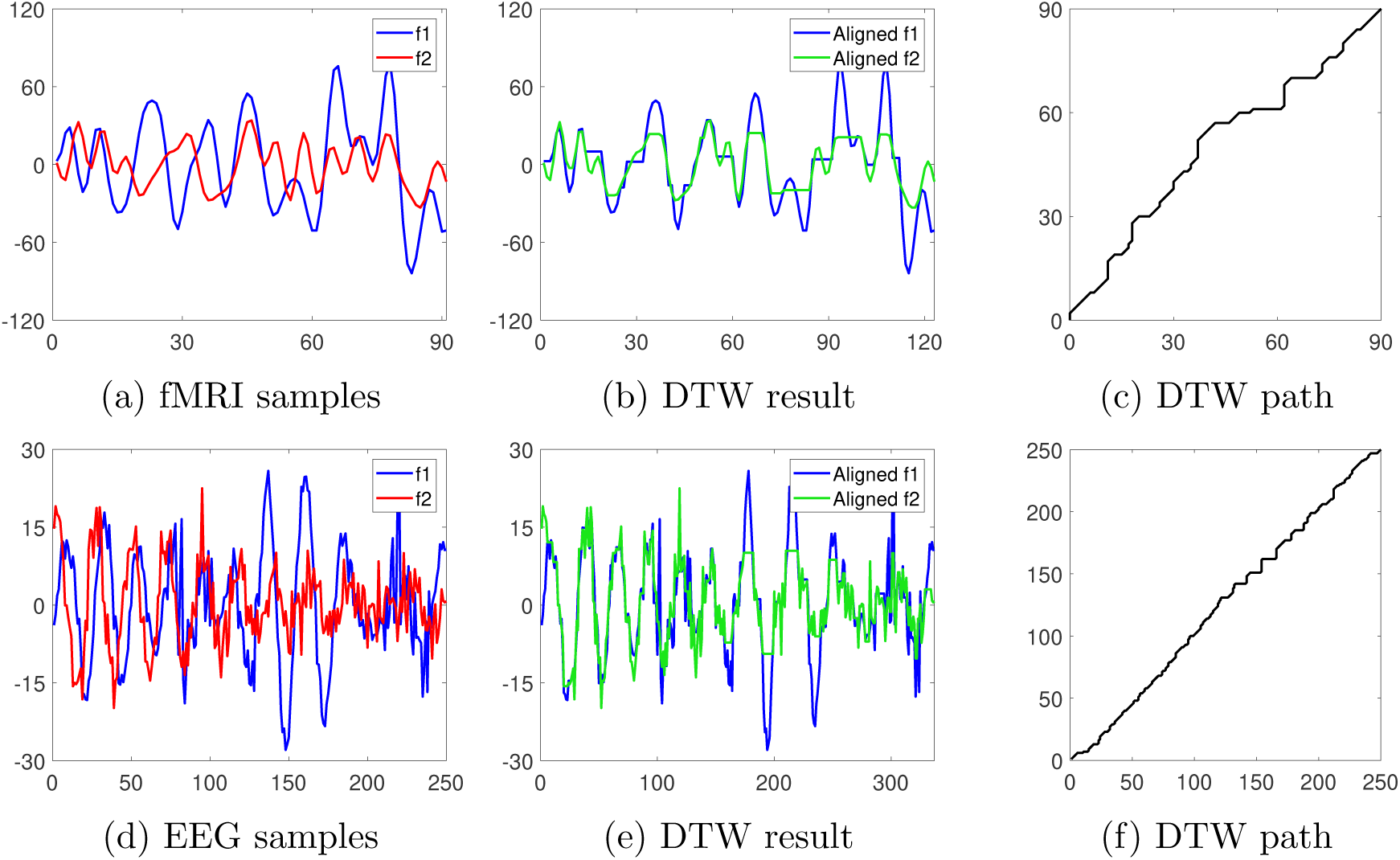
Illustration of the DTW method on real data. **(a)** One pair of fMRI signals *f*_1_ and *f*_2_, represented with 90 discrete sampling points. These signals are identical to those in Fig.7 (a); **(b)** Aligned signals via DTW; **(c)** Shortest path for the optimal DTW match; **(d)** One pair of EEG signals *f*_1_ and *f*_2_, represented with 250 discrete sampling points. These signals are identical to those in Fig.7 (d); **(e), (f)** Same as (b) and (c) except for the EEG signals.

### 3.2 Brain network via the FRR method

Recent study on brain network has been focusing on functional methods, where neural signals are treated as real-valued functions over a continuous time domain. A graph-based method, called functional additive semi-graphoid (FASG) model, was developed to build network using fMRI data from different brain areas (Li and Solea, 2018). This functional graphical model is based on the notion of Additive Conditional Independence (ACI) (Li et al., 2014), where random functions are assumed to reside in two level Hilbert spaces – {ℋ_*i*_}_*i*=1,…, *p*_ as the first level and a set of {ℳ_*i*_}_*i*=1,…, *p*_ RKHS as the second Hilbert spaces, where *p* denotes the number of vertices in the graph. These two levle spaces are connected by a set of positive definite mapping {*κ*_*i*_(·, ·)}_*i*=1,…, *p*_ from ℋ_*i*_ × ℋ_*i*_ → ℝ as kernel functions. One of the key benefits of this method is its nonlinear, non-Gaussian property. This is a more general and powerful than conventional Gaussian-based methods (Meinshausen et al., 2006; Qiao et al., 2018).

In practice, the mapping kernel is often taken in the following form: 

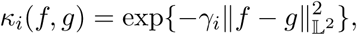

where *γ*_*i*_ *>* 0 is a scale parameter and the norm 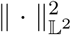 denotes the Euclidean 𝕃^2^ distance. In this study, we introduce the FRR as a pre-processing procedure to remove the phase variability among one area and then construct the graph with the aforementioned setting. We will compare the network results of three methods: 1) Euclidean distance on the original data, 2) The DTW distance, and 3) Euclidean distance on the aligned data via the FRR.

#### 3.2.1 Network analysis – fMRI data

We now apply the FASG model under three settings to the fMRI data set, described in Sec. 3.1.1. With 116 ROIs, there are 116 nodes in the network. The number of edges in a fully connected network is 116 *×* 115*/*2 = 6670. In this study, we construct brain networks by connecting edges with top 5% strongest additive dependence in the FASG method. That is, we will have 6670*×*0.05 = 334 edges in our estimated network.

In Figure 9, the constructed networks are shown for two groups by three settings, namely, {marijuana users, non-marijuana users}*×*{Euclidean distance, DTW distance, Euclidean distance after FRR alignment}. Visually speaking, the three networks in the upper row (for non-marijuana uses) look very similar. At a more detailed level, we can see high similarity between networks obtained by FRR and DTW, which are somewhat different from that by Euclidean distance. In contrast, the three networks in the lower row (for marijuana users) look very much different. These results indicate that the three methods can yield substantial differences in the marijuana user group.

**Figure 9:**
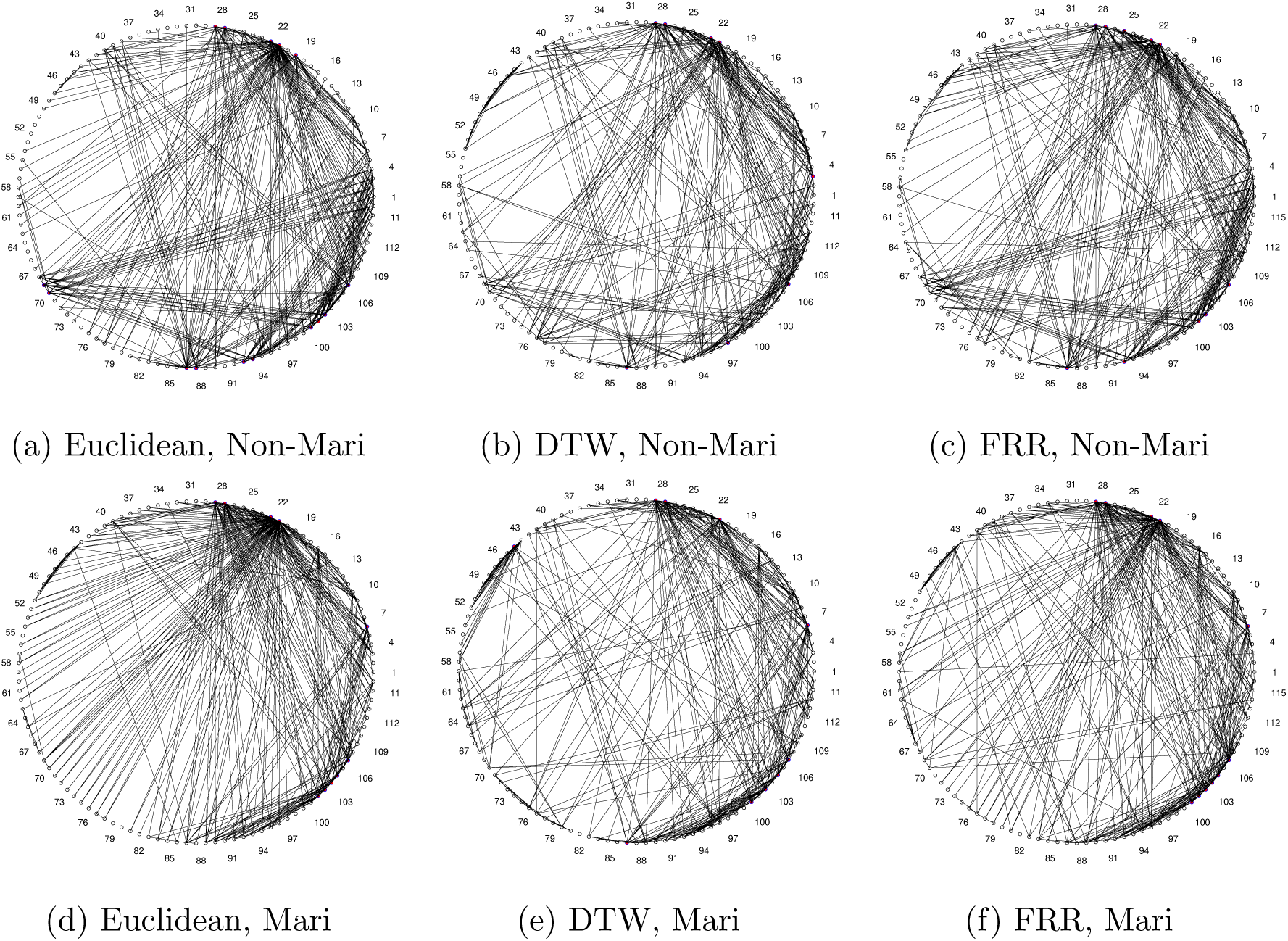
Brain networks constructed by the FASG model on the fMRI data. **(a)**: Network obtained by Euclidean distance for non-marijuana group; **(b)**: Network obtained by DTW distance for non-marijuana group; **(c)**: Network obtained by Euclidean distance after FRR alignment for non-marijuana group; **(d), (e), (f)**: Same as (a), (b), (c) except for the marijuana group.

For each method, we can also observe some difference between the non-marijuana-user group and marijuana-user group. We can assess whether the difference is statistically significant by a random permutation approach (Li and Solea, 2018). That is, we can check if the difference between the two groups is due to group variation or random variation among individuals. This is quantified by the difference of the covariance operators of the two groups. Here we use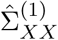and 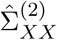 to denote the covariance operators of the non-marijuana-user and marijuana-user groups, respectively. We compute the difference 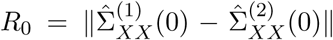 for the original group assignment, and 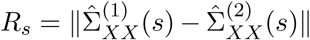, *s* = 1, …, 100, for 100 rounds of random permutation. In each round, we randomly assign 30 out of the total 60 as marijuana users, and the other 30 as non-marijuana users. Figure 10 shows the histograms of all *R*_*s*_ and the location of *R*_0_. We can see that under the DTW and FRR settings (after removing pairwise phase-lags), *R*_0_ for the true sample locates near the right end of all *R*_*s*_ produced by random permutations. This indicates that, for DTW and FRR, the difference between the two groups is statistically significant. In contrast, the *R*_0_ obtained by Euclidean distance is near the center of the distribution of *R*_*s*_, which implies we cannot tell if the detected difference obtained in Figure 9 is due to group variation or individual variation.

**Figure 10:**
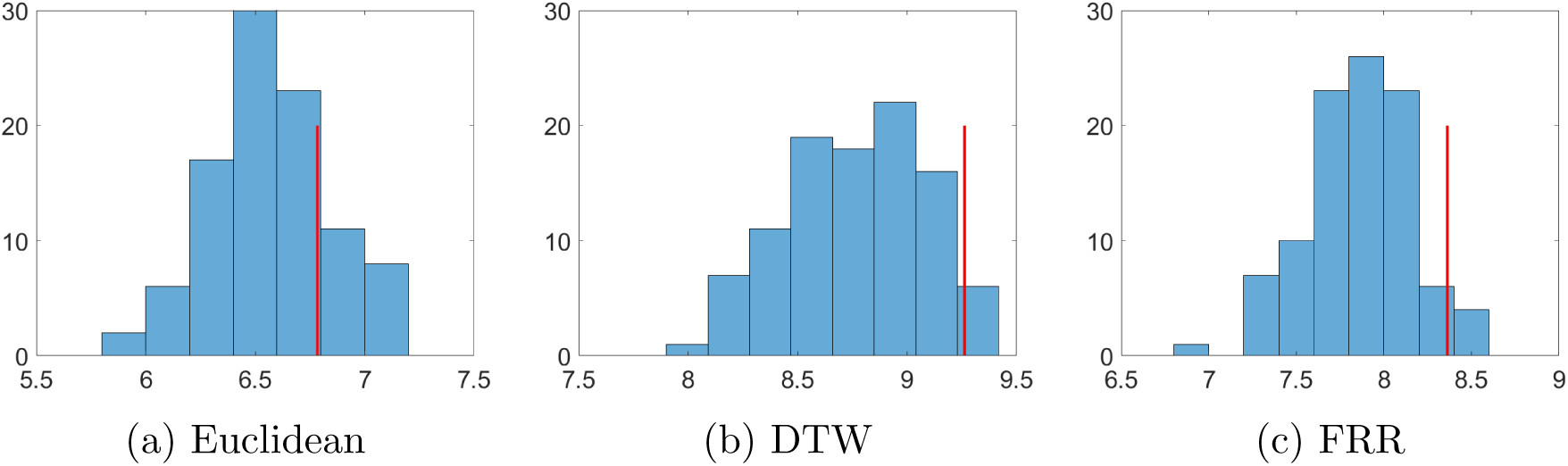
Histogram for covariance operator difference 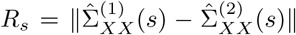, for *s* = 1, …, 100 randomly split samples and the position of *R*_0_ for true sample under different settings, where the red line indicates the position of *R*_0_: **(a)** Euclidean - *R*_0_ is larger than 79% of 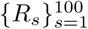; **(b)** DTW - 96%; **(c)** FRR - 97%.

To further investigate the difference between groups, we list the regions with more than 20 edges in Table 2. From this table, we can see that in each of the three method, the number of high-connectivity regions is always slightly higher (by 1) in the marijuana-user group. In particular, the

**Table 2:** Vertices with more than 20 edges

|  | Euclidean | DTW | FRR |
| --- | --- | --- | --- |
| Non-marijuana | 21, 22, 87 | 21, 22, 27, 28 | 21, 22, 27, 28, 87 |
| Marijuana | 21, 22, 27, 28 | 21, 27, 28, **103** | 21, 22, 27, 28, **105, 107** |

DTW and FRR models both identify high-connectivity regions in the cerebellum (103, 105 and 107) in the marijuana group. The extant literature indicates that long-term daily cannabis users showed an increase in cerebellar blood volume (Sneider et al., 2006). This is well captured by the DTW and FRR methods when the phase variability is removed in the given fMRI signals. In summary, we see more high-connectivity regions in the marijuana group compared to the non-marijuana group, and the increase in cerebellar blood volume is identified by removing phase variability using the DTW and FRR methods.

#### 3.2.2 Network analysis – EEG data

We also apply the FASG model under three settings to the EEG data set. With 32 electrodes, there are 32 nodes in the network. The number of edges in a fully connected network is 32 *×* 31*/*2 = 496. Same as the fMRI case, we construct brain networks by connecting edges with top 5% strongest additive dependence. That is, we will have 496 *×* 0.05 = 25 edges in each network.

The estimated networks using three distances methods on Day 1 and Day 4 are shown in Figure 11. Basically, there is no much difference between three networks for Day 1 (shown in the upper panels). All of them show many connections in the posterior area (17, 18, 19, 20), some connections in the central region (7, 8, 9, 11, 22, 24), and very few common edges ({(2, 3), (27, 31)}) in the frontal area. However, for Day 4, the network under Euclidean setting is very different from the other two. That is, high connectivity is only detected in the posterior area using the Euclidean distance, whereas the networks obtained by DTW and FRR are constructed alike – they both show high connections between the frontal area and the posterior area. It was found that tACS induces long-term increases in posterior-to-frontal alpha connectivity (Clancy et al., 2018). The networks indicate that this long-term enhancement effect is confirmed by the FASG graph model under the DTW and FRR settings (pairwise phase-lags are removed).

**Figure 11:**
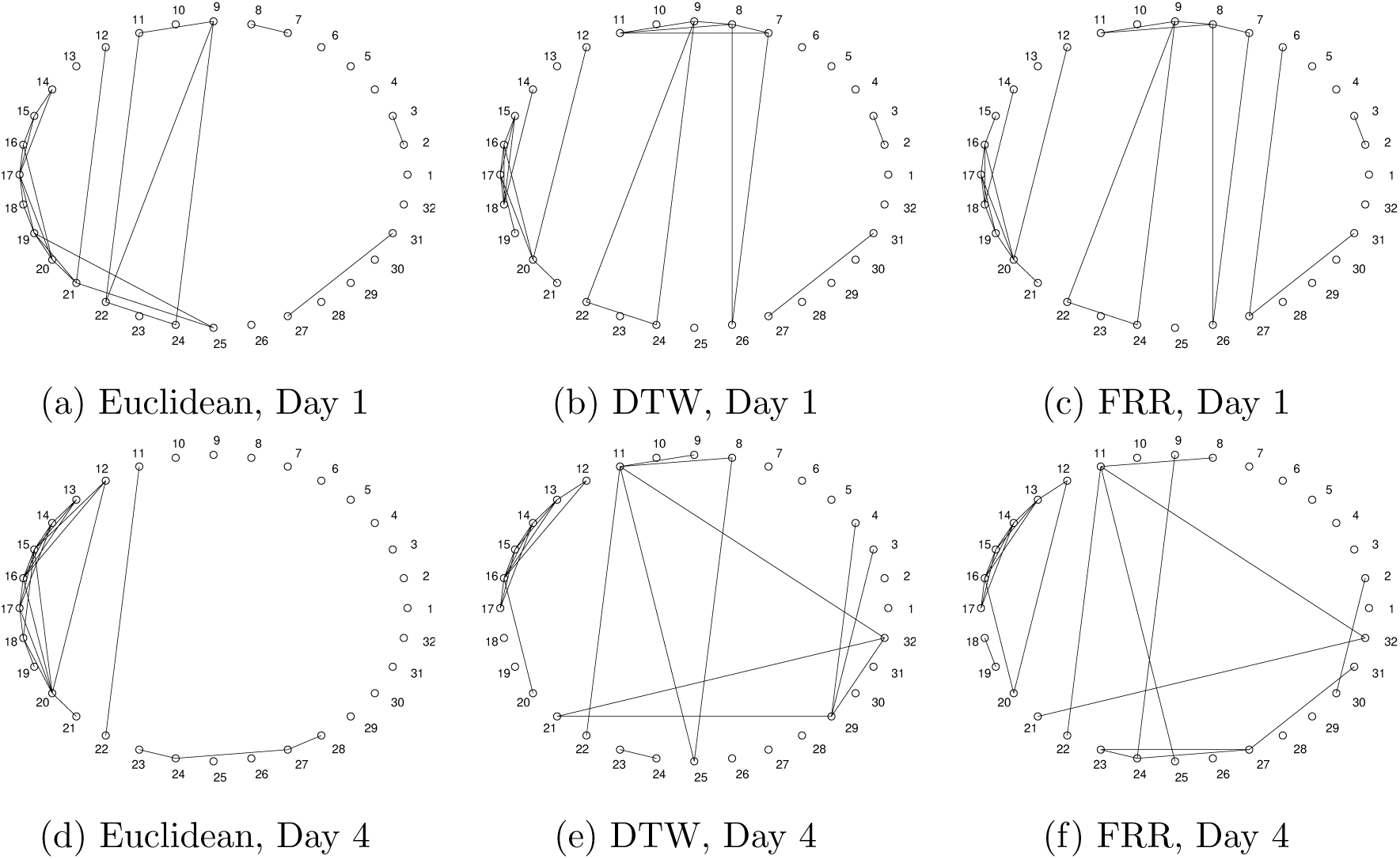
Brain networks constructed by the FASG model on the EEG data. **(a)**: Network obtained by Euclidean distance for Day 1; **(b)**: Network obtained by DTW distance for Day 1; **(c)**: Network obtained by Euclidean distance after FRR for Day 1; **(d), (e), (f)**: Same as (a), (b), (c) except for Day 4.

Similar to the result on the fMRI data, we also observe difference between Day 1 and Day 4 using each distance. To evaluate if the difference is significant, we again conduct the permutation-based resampling. Results are shown in Figure 12. It is found that only *R*_0_ for FRR locates at 97% (above the 95% significance threshold), whereas *R*_0_ for Euclidean and DTW is below the significance threshold. That is, the difference between two different days, reflecting the effect of tACS, can be effectively identified using the FRR method, whereas the Euclidean and DTW fail to detect the difference.

**Figure 12:**
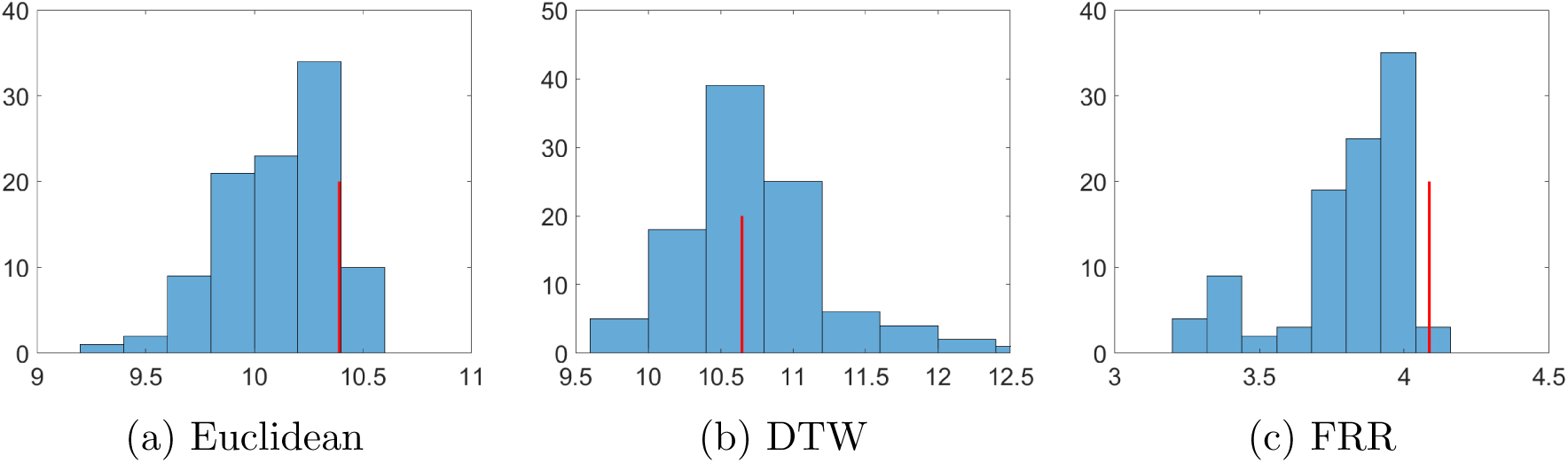
Histogram for covariance operator difference 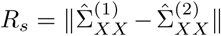, for *s* = 1, …, 100 randomly split samples and the position of *R*_0_ for true sample under different settings, and the red line indicates the position of *R*_0_: **(a)** Euclidean - *R*_0_ is larger than 89% of 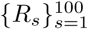; **(b)** DTW - 51%; **(c)** FRR - 97%.

## 4 Discussion

Phase variability is commonly observed in various neural signals such as EEG and fMRI. Modeling and analyzing those signals has been a major challenge, especially when the variability is nonstationary and nonlinear. In this article, we propose to adopt the Fisher-Rao registration (FRR) framework to deal with this problem. We systematically compare our method to the well-known DTW method in three aspects - basic framework, mathematical properties, and computational efficiency. The comparative result clearly demonstrates the superior performance of the FRR method. We then apply the FRR method to brain network problems using an fMRI and an EEG dataset. It is found that the FRR method can successfully identify difference between states/groups in both datasets, whereas the DTW fails the task in the EEG dataset.

The FRR method shows its potentials in analyzing neural signals with phase variability. However, so far we have only tested the method with one state-of-the-art nonparametric graph model in the brain network analysis. We will explore other network models and test the method using more extensive data. We will also investigate our framework to study the problem of phase synchrony between brain regions (Lachaux et al., 1999). Finally, the current study focuses on continuous signals such as fMRI and EEG. We will explore possibilities to examine latency in discrete spike trains between neuronal units.

### A The DTW Algorithm

#### Algorithm 1 Dynamic Time Warping

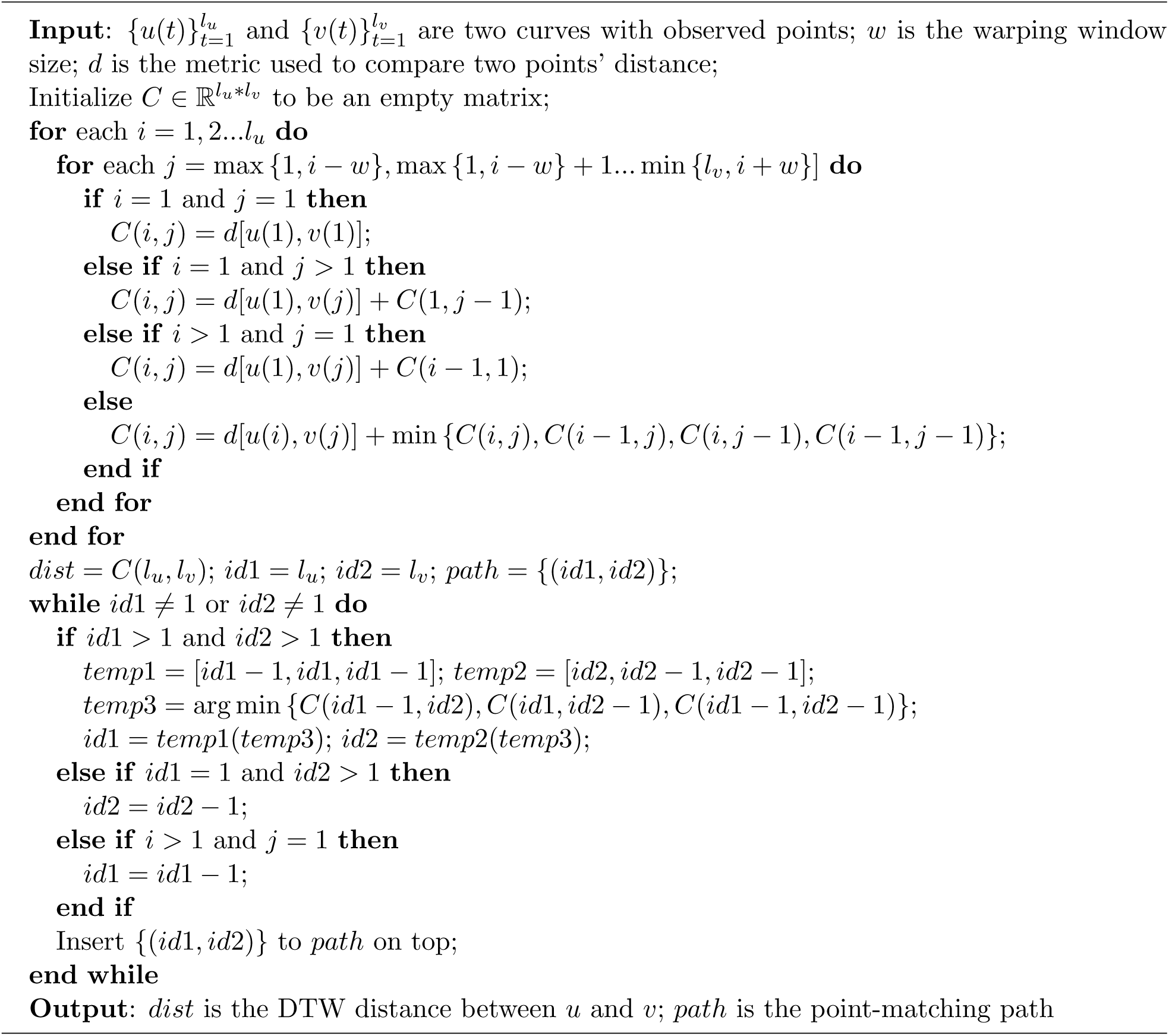

### B The Template Estimation Algorithm

The optimal template can be computed under the FRR method by:

1. Compute the SRVFs for the given curves: *q*_1_, *q*_2_…*q*_*n*_;
2. Iteratively: find the mean of the SRVFs *µ*; match each of the SRVFs to *µ* and replace the original ones; do until convergence to obtain 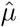, the estimated mean in amplitude (the Karcher mean);
3. Match each of the SRVFs to 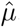 to obtain the estimated phase 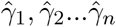;
4. Since the SRVF of a time warping function is in 𝕊^*∞*^, the mean of 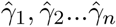 can be computed by the mean of their SRVFs (the Karcher mean), 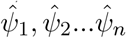, on the infinite-dimension ball, using the shooting vectors’ average and exponential mapping. In this way, the estimated mean in phase, 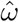, can be obtained;
5. Combining the average in amplitude and phase, one can obtain the optimal template:

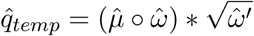

## Author Contributions

W.Z. designed the proposed framework, conducted computational analysis, and wrote the paper. Z.X. assisted with data analysis and wrote the paper. W.L. provided experimental data, interpretd analysis result, and wrote the paper. W.W. designed the proposed framework, supervised data analysis, and wrote the paper.

## Footnotes

1 https://github.com/WeilongZ/FRR-vs-DTW.git

2 http://fcon_1000.projects.nitrc.org/indi/ACPI/html/index.html

## References

Allen, E. A., Damaraju, E., Plis, S. M., Erhardt, E. B., Eichele, T., and Calhoun, V. D. (2014). Tracking whole-brain connectivity dynamics in the resting state. Cerebral cortex, 24(3):663–676.

Allen, E. A., Damaraju, E., Plis, S. M., et al. (2012). Tracking whole-brain connectivity dynamics in the resting state. Cerebral Cortex, 24(3):P663–676.

Brookes, M. J., Hale, J. R., Zumer, J. M., et al. (2011). Measuring functional connectivity using meg: Methodology and comparison with fcmri. NeuroImage, 56(3):P1082–1104.

Chang, C. and Glover, G. H. (2010). Time–frequency dynamics of resting-state brain connectivity measured with fmri. NeuroImage, 50(1):P81–98.

Chen, J. E., Chang, C., Greicius, M. D., and Glovera, G. H. (2015). Introducing co-activation pattern metrics to quantify spontaneous brain network dynamics. NeuroImage, 111:P476–488.

Clancy, K. J., Baisley, S. K., Albizu, A., Kartvelishvili, N., Ding, M., and Li, W. (2018). Lasting connectivity increase and anxiety reduction via transcranial alternating current stimulation. Social cognitive and affective neuroscience, 13(12):1305–1316.

Craddock, R. C., James, G. A., Holtzheimer III, P. E., Hu, X. P., and Mayberg, H. S. (2012). A whole brain fmri atlas generated via spatially constrained spectral clustering. Human brain mapping, 33(8):1914–1928.

de Pasquale, F., Penna, S. D., Snyder, A. Z., et al. (2010). Temporal dynamics of spontaneous meg activity in brain networks. PNAS, 107(13):P6040–6045.

Dinov, M., Lorenz, R., Scott, G., Sharp, D. J., Fagerholm, E. D., and Leech, R. (2016). Novel modeling of task vs. rest brain state predictability using a dynamic time warping spectrum: comparisons and contrasts with other standard measures of brain dynamics. Frontiers in computational neuroscience, 10:46.

Fries, P. (2005). A mechanism for cognitive dynamics: neuronal communication through neuronal coherence. Trends in Cognitive Sciences, 9(10):P474–480.

Friston, K. J., Buechel, C., Fink, G. R., et al. (1997). Psychophysiological and modulatory interactions in neuroimaging. NeuroImage, 6(3):P218–229.

Gupta, L., Molfese, D. L., Tammana, R., and Simos, P. G. (1996). Nonlinear alignment and averaging for estimating the evoked potential. IEEE Transactions on Biomedical Engineering, 43(4):P348–356.

Handwerker, D. A., Gonzalez-Castillo, J., D’Esposito, M., and Bandettini, P. A. (2012). The continuing challenge of understanding and modeling hemodynamic variation in fmri. NeuroImage, 62(2):P1017–1023.

Huang, H.-C. and Jansen, B. (1985). Eeg waveform analysis by means of dynamic time-warping. International journal of bio-medical computing, 17(2):135–144.

Hurtado, J. E. (2004). An examination of methods for approximating implicit limit state functions from the viewpoint of statistical learning theory. Structural Safety, 26(3):P271–293.

Jones, D. T., Vemuri, P., Murphy, M. C., et al. (2012). Non-stationarity in the “resting brain’s” modular architecture. PLOS One, 380(9845):P899–907.

Kamiński, M., Ding, M., Truccolo, W. A., and Bressler, S. L. (2001). Evaluating causal relations in neural systems: Granger causality, directed transfer function and statistical assessment of significance. Biological Cybernetics, 85(2):P145–157.

Karamzadeh, N., Medvedev, A., Azari, A., et al. (2013). Capturing dynamic patterns of task-based functional connectivity with eeg. NeuroImage, 66:P311–317.

Keilholz, S. D. (2014). The neural basis of time-varying resting-state functional connectivity. Brain Connectivity, 4(10):P769–779.

Kiviniemi, V., Vire, T., Remes, J., et al. (2011). A sliding time-window ica reveals spatial variability of the default mode network in time. Brain Connectivity, 1(4):P339–347.

Lachaux, J., Rodriguez, E., Martinerie, J., and Varela, F. J. (1999). Measuring phase synchrony in brain signals. Human Brain Mapping, 8(4):P194–208.

Lawlor, P. N., Perich, M. G., Miller, L. E., and Kording, K. P. (2018). Linear-nonlinear-time-warp-poisson models of neural activity. Journal of computational neuroscience, 45(3):173–191.

Li, B., Chun, H., and Zhao, H. (2014). On an additive semigraphoid model for statistical networks with application to pathway analysis. Journal of the American Statistical Association, 109(507):1188–1204.

Li, B. and Solea, E. (2018). A nonparametric graphical model for functional data with application to brain networks based on fmri. Journal of the American Statistical Association, 113(524):1637–1655.

Li, Z., Liu, H., and Liao, X. (2015). Dynamic functional connectivity revealed by resting-state functional near-infrared spectroscopy. Biomedical Optics Express, 6(7):P2337–2352.

Liu, X. and Duyn, J. H. (2013). Time-varying functional network information extracted from brief instances of spontaneous brain activity. PNAS, 110(11):P4392–4397.

Meinshausen, N., Bühlmann, P., et al. (2006). High-dimensional graphs and variable selection with the lasso. The annals of statistics, 34(3):1436–1462.

Meszlényi, R. J., Hermann, P., Buza, K., et al. (2017). Resting state fmri functional connectivity analysis using dynamic time warping. Frontiers in Neuroscience, 11.

Mørup, M., Hansen, L. K., and Arnfred, S. M. (2008). Algorithms for sparse nonnegative tucker decompositions. Neural Computation, 20(8):P2112–2131.

Qiao, X., Guo, S., and James, G. M. (2018). Functional graphical models. Journal of the American Statistical Association, pages 1–12.

Sakoe, H. and Chiba, S. (1978). Dynamic programming algorithm optimization for spoken word recognition. IEEE transactions on acoustics, speech, and signal processing, 26(1):43–49.

Sakoğlu, Ü., Pearlson, G. D., Kiehl, K. A., et al. (2010). A method for evaluating dynamic functional network connectivity and task-modulation: application to schizophrenia. Magnetic Resonance Materials in Physics, Biology and Medicine, 23(5-6):P351–366.

Singer, W. (1999). Neuronal synchrony: A versatile code for the definition of relations. Neuron, 24(1):P49–65.

Singer, W. and Gray, C. M. (1995). Visual feature integration and the temporal correlation hypothesis. Annual Review of Neuroscience, 18(Singer):P555–586.

Smith, S. M. (2012). The future of fmri connectivity. NeuroImage, 62(2):P1257–1266.

Sneider, J. T., Pope Jr, H. G., Silveri, M. M., Simpson, N. S., Gruber, S. A., and Yurgelun-Todd, D. A. (2006). Altered regional blood volume in chronic cannabis smokers. Experimental and clinical psychopharmacology, 14(4):422.

Srivastava, A., Wu, W., Kurtek, S., Klassen, E., and Marron, J. (2011). Registration of functional data using fisher-rao metric. arXiv preprint 1103.3817.

Tononi, G., Edelman, G. M., and Sporns, O. (1998). Complexity and coherency: integrating information in the brain. Trends in Congitive Sciences, 2(12):P474–484.

Tucker, J. D., Wu, W., and Srivastava, A. (2013). Generative models for functional data using phase and amplitude separation. Computational Statistics & Data Analysis, 61:50–66.

Varela, F., Lachaux, J.-P., Rodriguez, E., and Martinerie, J. (2001). The brainweb: Phase synchronization and large-scale integration. Nature Reviews Neurosciencevolume, 2(4):P229–239.

Vinck, M., Oostenveld, R., van Wingerden, M., et al. (2011). An improved index of phase synchro-nization for electrophysilolgical data in the presence of volume-conduction, noise and sample-size bias. NeuroImage, 55(4):1548–1565.

Williams, A. H., Poole, B., Maheswaranathan, N., Dhawale, A. K., Fisher, T., Wilson, C. D., Brann, D. H., Trautmann, E. M., Ryu, S., Shusterman, R., et al. (2020). Discovering precise temporal patterns in large-scale neural recordings through robust and interpretable time warping. Neuron, 105(2):246–259.

Zhang, J.-T. (2013). Analysis of variance for functional data. CRC Press.

